# Timing and clustering co-occurring genome amplifications in cancers

**DOI:** 10.1101/2025.03.31.646456

**Authors:** Sara Cocomello, Giovanni Santacatterina, Riccardo Bergamin, Alice Antonello, Giulio Caravagna

## Abstract

Clonal evolution in cancer is driven by genomic alterations that accumulate over time, shaping tumour progression, therapy resistance, and metastasis. Among these somatic events, genomic amplifications are a broad class of copy number alterations (CNAs) that can be mathematically timed (i.e., mapped to an abstract timeline). Existing methods successfully order amplifications in time but fail to understand their co-occurrence patterns. This limitation makes it harder to understand abrupt shifts of clonal and selection dynamics possibly linked to clones that acquire profound mutant genotypes and hold the potential to establish a novel evolutionary lineage. Here, we introduce `TickTack`, a hierarchical Bayesian mixture model for reconstructing the temporal order of copy number amplifications across the genome while simultaneously detecting co-occurrent events, offering a more comprehensive view of tumour evolutionary dynamics. This new model allows us to determine whether copy number amplifications accumulate gradually over multiple generations or occur in rapid succession within short time frames, providing deeper insights into genomic instability and tumor progression beyond traditional linear models. We validated our approach with synthetic data under various uncertainty settings and against competing approaches. Applying TickTack to 2,777 samples from the Pan-Cancer Analysis of Whole Genomes (PCAWG) project, a comprehensive resource spanning 38 tumor types, we inferred the temporal order of copy number amplifications, identifying cancer-specific co-occurring events. Our analysis revealed associations between early chromosomal instability and key driver mutations (TP53, BRCA1/2) in Esophageal Adenocarcinoma and uncovered recurrent evolutionary trajectories shaped by focal and arm-level copy gains. These findings highlight the role of saltational evolution in tumorigenesis and provide insights into genomic instability with possible implications for prognosis and targeted therapies.

**Availability:** `tickTack` is available as an R package at https://caravagnalab.github.io/tickTack/ and the code to replicate the analysis is available from https://zenodo.org/records/14870458.

## 1 Introduction

The temporal dynamics of clonal evolution play a critical role in shaping the trajectory of cancer development [1, 2]. Understanding when specific genomic events occur within a tumour can provide invaluable insights into the mechanisms driving oncogenesis, therapy resistance, and metastasis [3]. Among the most significant of these events are copy number alterations (CNAs), which represent gains or losses of genomic segments that contribute to tumour heterogeneity and progression [4, 5].

Despite their importance, inferring the timing and co-occurrence of CNAs across the genome remains a considerable computational challenge, with modern methods primarily focusing on copy number amplifications (or copy gains) [6, 7]. Analyzing co-occurring amplifications can reveal crucial evolutionary events involving key tumour suppressor genes and oncogenes, including whole-genome doubling (WGD) events and “hopeful monsters”. Whereas WGD is a well-studied phenomenon in cancer [8–10], the hopeful monsters paradigm by Goldschmidt posits that a combination of simultaneous alterations – hereby copy gains – can lead to phenotypic changes substantial enough to overcome evolutionary bottlenecks and establish a new clonal population [11, 12]. By unraveling the interplay and timing between these genomic alterations, we can better reconstruct the evolutionary steps that promote tumorigenesis [13, 14].

Several computational methods have been developed to time CNAs [15–17]. MutationTimeR, first used to analyse the Pan-Cancer Analysis of Whole Genomes (PCAWG) cohort [18], calculates the relative timing of copy gains by comparing the number of point mutations at different Variant Allelic Frequency (VAF) within altered genomic segments [16]. The intuition is that mutations occurring before a copy gain on the amplified allele will be present on both the original and the amplified allele, hence will have higher VAF, while mutations sitting on the non-amplified allele or occurring after the amplification will be present on a single allele, hence will have a lower VAF. In other words, if we define the multiplicity of a mutation as the number of alleles on which the mutation is found, for copy gains we will have that mutations on amplified alleles have a multiplicity equal to 2, while mutations on the non-amplified allele or occurring after the copy gain will have multiplicity 1. This method assumes a constant mutation rate, cannot model complex multi-allele amplifications, or detect co-occurring events. Recently, AmplificationTimeR extended MutationTimeR to include more complex copy number gains, such as sequential amplifications and WGD [19]. AmplificationTimeR estimates the order of events for regions undergoing multiple amplifications, providing detailed insights into the temporal sequence of amplifications. Its application is, however, hindered by the computational complexity of modelling large-scale genomic events and the need for high-quality multiplicity data [20]. Moreover, like MutationTimeR, AmplificationTimeR cannot detect which amplifications happened together.

By treating genomic regions independently, both MutationTimeR and AmplificationTimeR overlook the interplay of co-occurring alterations. Addressing this limitation requires models to integrate co-occurrency into the inference process. In this study, we built on these principles to introduce `TickTack`, a hierarchical Bayesian mixture model designed to time copy gains and determine their co-occurrence patterns. We initially developed the method for single-segment inference, which we refer to as `TickTack` baseline, providing a foundation for modeling individual events before extending it to account for co-occurring alterations (`TickTack` full). Our model enables the genome-wide inference of copy gains that likely happened at the same time, offering a principled way to detect complex aneuploidy patterns like hopeful monsters (Fig.1a). due versioni una singolo segmento una clustering By applying TickTack to the PCAWG cohort spanning 32 cancer types, we demonstrated its ability to identify the relative order of events in tumor clonal evolution. Our analysis revealed cancer-specific co-occurring patterns, including significant associations between early chromosomal instability and key mutations (TP53, BRCA1/2) across multiple cancer types, as well as recurrent temporal sequences in both focal and arm-level copy gains. Notably, we found that TP53 in Esophageal Adenocarcinoma usually precedes CCND1, BRCA1, and ABL1 and that the 17p arm, where TP53 is located, consistently appears early in LIRI, preceding the amplification of 10q. These results highlight the potential of our approach to uncover the temporal structure of genomic instability and its relationship with driver mutations during tumorigenesis. Our work represents a significant advance in computational methods for cancer genomics, providing a framework for studying the ordered progression of copy number alterations and their connections to cancer-driving mutations.

**Fig. 1.**
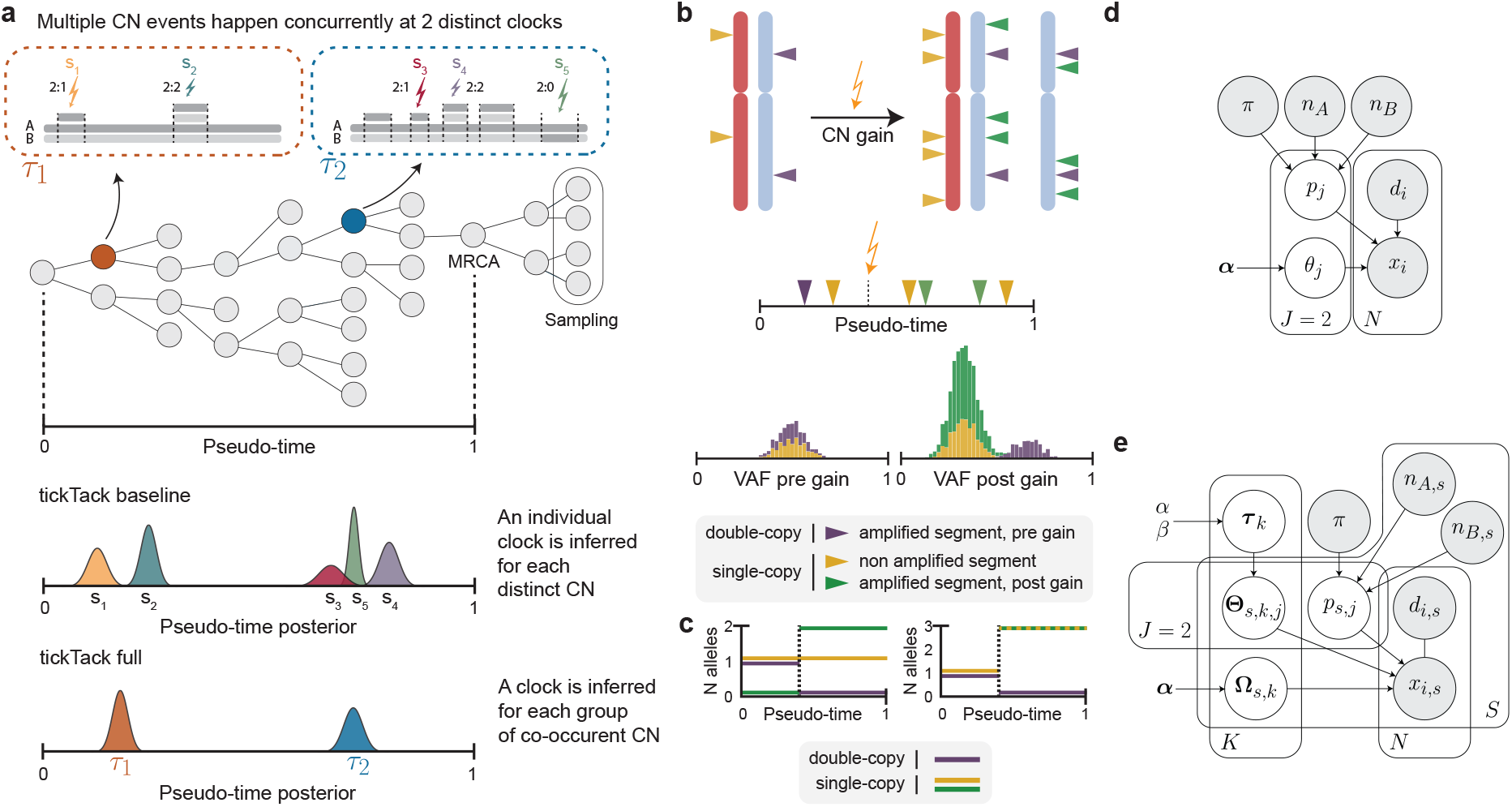
Timing genome amplifications in cancer: the **TickTack** approach. **(a)** Phylogenetic tree illustrating the clonal evolution of copy number gains, with 5 events occurring at two distinct clocks. The lower panels show results from the single- and multiple-segment inference approaches, highlighting differences between the two approaches. `TickTack` baseline estimates 5 different clocks for each affected genomic segment, while `TickTack`full is able to correctly assign each segment to one of the two biologically relevant clocks. **(b)** Copy number gains are allelic amplifications that affect the multiplicity of cancer mutations, which can be distinguished by when and where they arrive on the genome (allelic position). We can map the time of copy gains along a pseudo-time [0, 1] axis. Bimodal variant allele frequencies (VAF) position the time towards the beginning or end of the interval. **(c)** The number of alleles harboring mutations in single- and double-copy changes after a copy number gain. The case of trisomy (2:1) is depicted. Before the gain, 1 allele harbors mutations in double-copy, and 1 allele harbors mutations in single-copy. On the other hand, after the gain, all three alleles harbor mutations in single-copy. **d**,**e)** Probabilistic graphical models for timing copy number gains. (d) Single-segment model where each segment is timed independently using a binomial mixture model with weights *θ*_*j*_, which determine the amplification time *τ*. (e) Hierarchical model that allows multiple segments to share *K* distinct amplification times *τ*_*k*_ through segment-specific mixture weights **Ω**, enabling the detection of co-occurrent amplification events.

## 2 Model description

Timing genome amplification events can be done using Poisson processes [1] and several simplifying assumptions shared among most models. First, the mutation rate *ρ >* 0 is assumed constant over time and across segments, which might be inaccurate when patients receive mutagenic treatments or specific mutations lead to microsatellite instability [21]. Then, mutations are supposed to follow the infinite site model (i.e., each site in the genome is mutated only once), and the likelihood of acquiring a mutation on each chromosome copy is equal. Moreover, at the mechanistic level, bi-allelic gains are assumed to happen simultaneously, which might hold for whole genome doubling events that double chromosome counts simultaneously but might be generally inaccurate. In general, timing is only performed for clonal copy gains (i.e., copy gains present in all tumour cells, occurred before the sample’s most recent common ancestor, MRCA [17]). Finally, a further assumption specific to our model and MutationTimeR [16] is to time just simple amplifications, which include copy-neutral loss-of-heterozygosity (CNLOH) events, as well as gain of alleles that lead to trisomy or tetrasomy amplifications.

### 2.1 Timing segments independently

We start with a model to time, independently, a set of clonal copy number gains. Specifically, by reframing the ideas first introduced by Nik-Zainal et al.[22] within a Bayesian framework, we introduce `TickTack` baseline. The goal of the method is to time single amplification events and map each one to a pseudo-time 0 ≤ *τ* ≤ 1, an abstract representation of the lifespan of a cancer. A value of 0 will correspond to the date of birth of the cancerous population that we are studying, while 1 to the date on which the MRCA of the cells in the sample was born.

*Amplification clocks as mixture models* CNAs cause chromosome copy changes that, in turn, impact the VAFs of the somatic mutations that sit on the altered genome segments (Fig.1b). In the case of the clonal copy gains considered for our timing model and taking as input read count data, i.e. number of variant reads **x** and read depths **d**, the likelihood for the *i*^*th*^ mutation on the segment can be approximated as a distribution with two modes

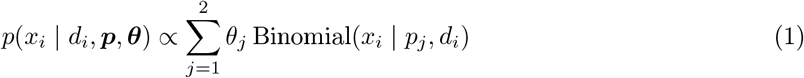

where ***θ*** = (*θ*_1_, *θ*_2_) are the binomial mixing proportions and ***p*** = (*p*_1_, *p*_2_) are the mixture modes: 0 ≤ *p*_2_ ≤ 1 is for mutations accumulated before the gain on the amplified segment, and therefore in double copy, and 0 ≤ *p*_1_ ≤ 1 is for mutations accumulated after the gain on any segment, or on the non-amplified segment, and therefore in single copy (Fig.1b). The expected values for the two modes depend on the particular conformation of the involved chromosomes: if the segment has *n*_*A*_ copies of the major allele, and *n*_*B*_ copies of the minor, for every *j* ∈ 1, 2

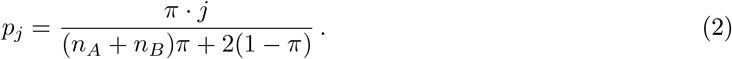

where 0 ≤ *π* ≤ 1 is the bulk sample purity. This distribution is just an approximation to the actual density because it neglects power-law distributed neutral mutations, whose frequency is below *p*_1_ by definition [23].

The key connection between the timing problem and the deconvolution is that the pseudo-time *τ* depends on the proportion of mutations in single-copy (*θ*_1_) and double-copy (*θ*_2_) states. These proportions can approximate the mutations counts, i.e *θ*_*i*_ ~ *N*_*i*_*/N*, with *N* being the total number of observed mutations. Hence, the full relationship between *τ, θ*_*i*_, and *N*_*i*_ can be understood by defining two molecular clocks to model the accumulation of mutations on the target segment. If we consider a segment of length *L*, chromosome copies *a*, cell division rate *ω >* 0 (cell doublings), and mutation rate *ρ >* 0 (mutations per chromosome, per cell doubling), we can write the time-inhomogeneous Poisson process

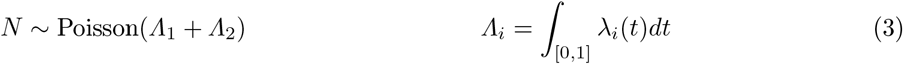

with intensity *λ*_*i*_(*t*), *i* ∈ 1, 2, that depends on *L, a, ω* and *µ*. Note that *Λ*_1_ + *Λ*_2_ is also the expectation for *N*. This clock is based on the closure of Poisson distribution under the sum operator, so that

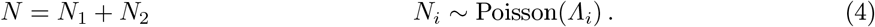

These two clocks can be defined considering that *N*_2_ mutations arrived on the genome before the amplification and were then amplified. Their rate is

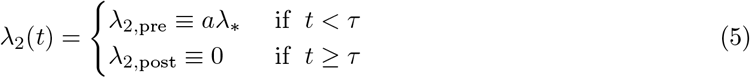

where *λ*_*_ = *Lωρ, a* = 1 for amplifications that lead to CNLOH and trisomy states, and *a* = 2 for GD events (Fig.1c). Note that the rate is 0 after *τ*, because no other amplifications are assumed to happen.

Instead, for *N*_1_ mutations arrived either after amplification or on the non-amplified allele

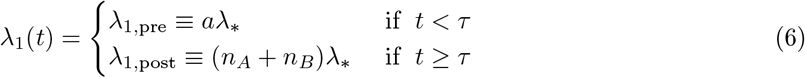

where *n*_*A*_ + *n*_*B*_ is the total segment ploidy after the amplification (2 for CNLOH, 3 for trisomy and 4 for GD) (Fig.1d). Instead, *a* = 0 for CNLOH because one allele is lost, *a* = 1 for trisomy and *a* = 0 for GD because both alleles are amplified.

We, therefore, obtain the time-homogenous piece-wise linear intensity

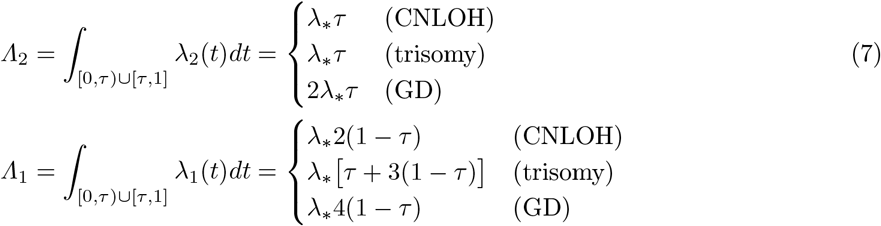

from which *N*_*i*_ ~ *Λ*_*i*_, *i* ∈ 1, 2 follows from the property of a Poisson process. For a specific copy number, one can, therefore, obtain *τ* by solving a linear system, obtaining

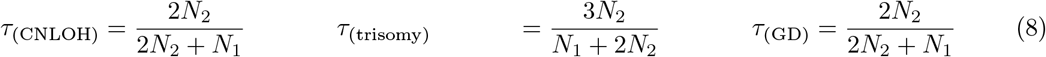

#### Likelihood

The overall likelihood for read count data from a segment affected by a simple copy gain is in Eq.(1). For this model, the prior on the mixture weights ***θ*** is ***θ*** ~ Dir(***α***) as we want them to sum to 1, being the probability of a mutation *i* to belong to the first or the second binomial component, setting *α* = 2 to allow for a quite sparse Dirichlet distribution. Therefore, the posterior for the parameters of interest will be

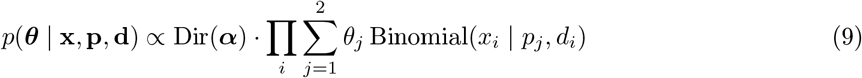

Once the posterior distribution over ***θ*** has been obtained, a posterior over *τ* is computed using Eq.(8), where *N*_*i*_ is replaced by *θ*_*i*_.

### 2.2 Detecting co-occurrent amplification events

Biological evidence suggests that multiple CNAs can arise from a single molecular event, resulting in near-simultaneous changes across different genome regions [6, 7]. This feature cannot be captured by assuming the independence of the timed segments. For this reason, we developed a hierarchical model that better reflects the underlying biology: rather than treating all *S* observed copy gains as independent, we propose that they arise from *K* distinct clocks, where *K* is typically much smaller than *S*. These clocks are macro-events representing distinct time points at which multiple genomic segments may have been simultaneously affected by copy number changes (Fig.1d).

#### Hierarchical model formulation

We extend our previous single-segment model into a hierarchical framework through a mixture-of-mixtures approach [24, 25]. questo costituisce il modello che chiameremo full ticktack. The model is augmented as follows: *i*) at the lower level, we still model each segment’s mutations using a mixture of two binomial distributions with parameters *θ*_1_ and *θ*_2_, *ii*) at the higher level, instead, we introduce a mixture component that allows segments to share *τ* values, and introduce a discrete set of *K* possible amplification times *τ*_1_, …, *τ*_*k*_. In this model, these shared timings induce dependencies between segments through their binomial parameters ***θ***, as segments with common *τ* must exhibit consistent distributions of mutations peaks, following Eq.(8). This hierarchical structure captures both intra-segment variability (through the binomial mixtures) and the inter-segment dependencies (through shared *τ* components). This constitutes the model we refer to as `TickTack` full. The complete model structure is visualized in (Fig.1e).

Similarly to the formulation presented in the previous section, the hierarchical model still employs the number of variant reads **x**, read depths **d**, and mixture modes **p**, as in Eq.(1-2). However, observations are now indexed by segment *s*, as we consider multiple segments simultaneously rather than treating them as independent. The key difference with the previous model is how the clocks are inferred. Previously, a single value of *τ* was estimated via Eq.(8) after determining the binomial mixture vector ***θ***. In contrast, we now infer a clock vector ***τ*** ∈ ℝ^*K*^. This vector, by the inverse of Eq.(8), generates a matrix ***Θ*** ∈ **R**^*S*×*K*×2^, representing binomial weights for single- and double-copy states across segments *s* and components *k*. Additionally, a weighting matrix **Ω** ∈ R^*S*×*K*^ is introduced. This matrix quantifies the degree to which each segment belongs to a particular component and is used for segment-to-cluster assignment. Thus, for the *i*^*th*^ mutation in segment *s*, the probability of observing variant reads *x*_*i,s*_ given read depth *d*_*i,s*_ and expected mixture modes **p**_*s*_ is

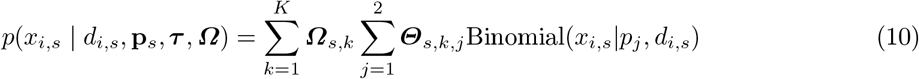

where **Ω**_*s,k*_ and ***Θ***_*s,k,j*_ represents mixing proportions for clock *k* and binomial weights, respectively. We stress again that the matrix ***Θ*** is a function of the clock vector ***τ***, which ultimately is our parameter of interest. For this model, we define the prior distributions

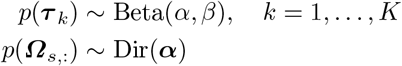

The Beta prior with hyper-parameters *α* and *β* is chosen such that the pseudo-time values are in the interval [0, 1] and the probability mass cover the interval in a quite homogeneous way. The Dirichlet distribution with concentration hyperparameters ***α*** is 5*/K* so that each cluster has a reasonable probability mass while still allowing the posterior to adjust based on the data. This avoids degenerate solutions where most of the weight is concentrated in just one or a few clusters, thereby maintaining flexibility in inference. Therefore, the full posterior is given by

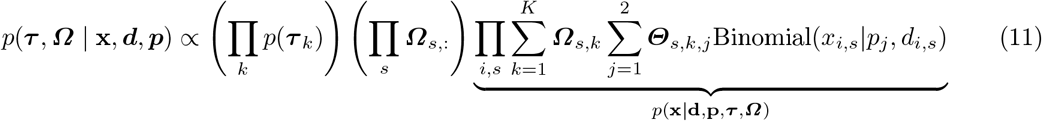

In this hierarchical clustering model, each segment is assigned to one of the *K* clocks based on the model likelihood. Once the posterior distributions of the parameters have been inferred, segment *s* is assigned to the most probable cluster by maximizing **Ω**_*s*,:_, i.e. *k*_*s*_ = arg max_*k*_ **Ω**_*s*,:_.

#### Determining the optimal number of clusters of amplifications

The method developed in this work is parametric: inference is performed iteratively with varying components, depending on the number of observed events, to optimize *K*. To balance model accuracy and complexity (number of clusters of amplifications), we adopted a model selection procedure based on the Akaike Information Criterion (AIC) [26] score, which was chosen by assessing the performance of various model selection criteria (e.g. BIC, LOO [27, 28]) on simulated data (Supplementary Fig.S1). AIC penalizes overly complex models (i.e., those with many components), while retaining a good fit to the data. In our case, its formula can be written as

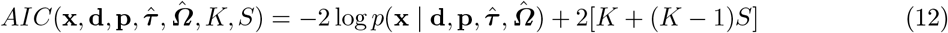

where the first term comes from Eq.(11) and the second term represents the number of parameters learned considering *S* segments and *K* clocks. Unlike other approaches, AIC penalizes the number of parameters by a constant factor of 2, regardless of the number of data points. For hierarchical models, AIC frequently suggests models with more components than alternative approaches [29].

#### tickTack implementation

We developed two probabilistic models for estimating the posterior distribution of `TickTack` parameters using Stan, a popular probabilistic programming for Bayesian inference that is wrapped in R[30]. Stan offers flexible modelling capabilities and supports both Markov Chain Monte Carlo (MCMC) and Variational Inference (VI), allowing a trade-off between statistical accuracy and computational efficiency.

In the first model (baseline `TickTack`, Section 2.1, (Fig.1e)), the posterior distribution of the key timing parameter *τ* is inferred from ***θ*** using Eq. (8). The parameters *θ*_1_ and *θ*_2_ are estimated via MCMC. This approach ensures robust inference by capturing the underlying data variability and dependencies but can be computationally demanding.

In contrast, the second model (full `TickTack`, Section 2.2, (Fig.1e)) adopts VI with automatic differentiation to infer the parameters ***τ*** and **Ω**, which respectively model the pseudotimes at which the events are distributed across the genome and segment-to-cluster assignments. VI is chosen for its scalability, particularly in scenarios where inference needs to be repeated multiple times to perform model selection for the number of clocks.

## 3 Validation on synthetic data and comparison with competing methods

### Synthetic data generation

To avoid any biases, we generated synthetic data following the AmplificationTimeR approach [19]. Cluster timing parameters (*τ*) were sampled from a uniform distribution [0.01, 0.99], ensuring a minimum pseudotime separation of 0.1 between the clocks generating amplification clusters. Each segment was assigned to a cluster, with *τ* perturbed by a 0-mean Gaussian noise (*σ* = 0.01). Using these values and mutation equations (7), expected mutation counts were computed for single and double-copy states, assigned proportionally within segments, and sampled from a multinomial distribution. Read counts were simulated using a binomial distribution, considering sequencing coverage and segment peak copy number.

To evaluate performance, we simulated various biological and computational scenarios by varying clocks(1–5), total number of amplification events (5–20), tumour purity (0.3–0.9), sequencing coverage (20–100), and mutations per segment (10–100). These scenarios represent a broad set of experimental and data settings, including whole-exome assays with few mutations, or very low-resolution data with low-purity samples. Each configuration was independently assessed and, for statistical reliability, we repeated each inference task 20 times to evaluate consistency and compare performance against alternative methods.

### Performance

We augmented MutationTimeR, AmplificationTimeR, and baseline `TickTack`(Section2.1) with a post-inference heuristic cluster step to allow a fair comparison against our full `TickTack`model (Section2.2). Since both MutationTimeR and AmplificationTimeR provided point estimates with confidence intervals (via bootstrap resampling) we applied hierarchical clustering [31] using a distance matrix based on confidence interval overlap, since the existing methods do not offer a way to cluster events for individual patients:

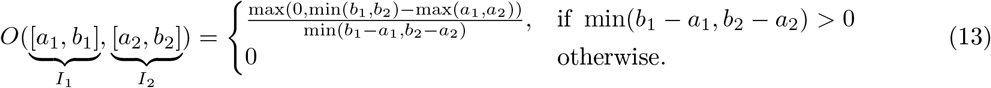

where *I*_1_ and *I*_2_ are the intervals considered. In contrast, for baseline `TickTack`, we used the full posterior distributions inferred via MCMC to compute their discrete Wasserstein [32] distance, a more informative distance metric between distributions

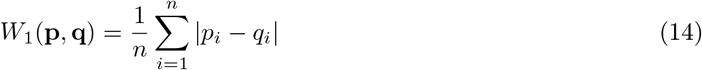

where **q** and **p** are two sorted vectors of posterior samples. We then constructed distance matrices and clustered segments using Ward’s method [33], determining the optimal number of clusters via Silhouette score[34]. Clustering performance was assessed by computing the rand index (RI)[35] between the groundtruth clustering (from simulations) and the inferred clusters, allowing us to establish the accuracy of each method in recovering the true evolutionary structure.

`TickTack` outperformed clustering, and performed well in approximating the real pseudotimes, particularly in scenarios with limited available data. In particular, `TickTack` achieved a median RI score of 0.88 across all simulations, surpassing alternative approaches significantly (median score 0.47; Fig.2a). The performance naturally depends on simulation complexity; for instance, in simulations with a single clock, `TickTack` attained a perfect median RI of 1. Even in more complex cases with five clocks, the score remained high at 0.84, demonstrating strong clustering capabilities. Interestingly, the second-best model is `TickTack` baseline, which performed slightly better than the competitors. We believe that this is linked with the clustering heuristic, which is based on posteriors that provide a richer representation of uncertainty and structure in the data.

**Fig. 2.**
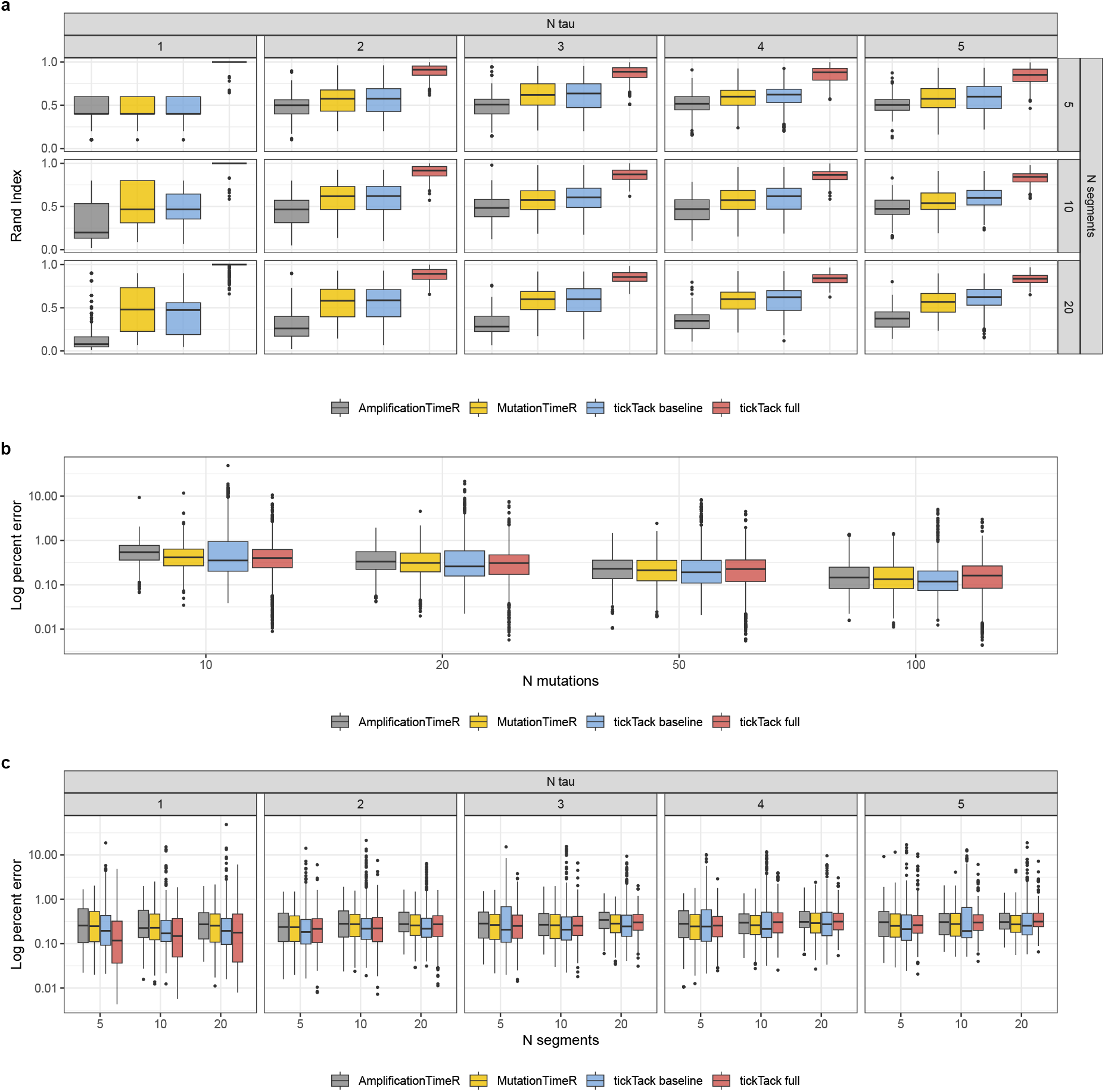
Comparison of timing inference methods on simulated data. **(a)** Rand Index measuring clustering accuracy. `TickTack` exhibits distinct performance, effectively grouping temporally close events compared to alternative approaches. **(b)** Percent errors of inferred clocks (against ground truth) for various number of mutations per segment (from few mutations, as in whole-exome, to many mutations, as in whole-genome assays). All methods perform better for richer datasets, with similar performance. **(c)** Percent errors of inferred clocks (against ground truth) across different numbers of segments and clocks (N tau). `TickTack` performances depend on the complexity of the simulation. Other methods’ performances are unaffected as performing inference independently per segment.

Considering the percent error of inferred clocks (against ground truth), `TickTack` improved timing accuracy, particularly in data-limited scenarios where the number of mutations per segment is low. In a whole-exome alike scenario (*N <* 10), where estimation uncertainty is high, `TickTack` achieved 0.23% lower error than AmplificationTimeR while performing comparably to MutationTimeR (0.40% vs. 0.41% median percentage error; Fig.2b). As expected, all methods improved with increasing mutation counts, with competitors’ errors dropping to 0.13%. In this regime, the advantage of `TickTack` for co-occurrent events diminished but remained competitive (0.16% median error). In contrast, `TickTack` baseline benefit from the additional data, improving to 0.12%. Unlike methods that infer timing independently per segment and remain largely unaffected by simulation complexity, `TickTack`’s performance adapts to the number of genomic segments and clocks (Fig.2c). Its error is lowest when fewer clocks are present (*τ* = 1, error = 0.14%), outperforming other methods (median error = 0.25%) and increasing slightly as complexity rises, though it remains comparable to competitors (0.29% vs. 0.30%). Overall, `TickTack` baseline consistently achieved the lowest error (0.21% median), followed by `TickTack` (0.26%), which performed on par with MutationTimeR and outperformed AmplificationTimeR (0.29%).

## 4 Timing of 1272 WGS samples across 32 cancer types

We applied `TickTack` to the PCAWG cohort, which includes 2,777 WGS samples spanning 47 tumor types. To refine our analysis, we selected 1,272 samples across 32 cancer types with a median of ≥ 10 mutations per segment. Ovarian cancers (OV) had the highest number of segments on average (~ 20 per sample), and diffuse large B-cell lymphoma (DLBC) the lowest (~ 1 per sample), consistent with known differences in genomic instability patterns [36]. The most prevalent copy gains (Supp.Figs.2-3) we timed was trisomy amplification (2:1), followed by tetrasomy (2:2) and copy-neutral loss of heterozygosity (CNLOH, 2:0), consistent with earlier work [20]

In Fig.3 we show the timing for two paradigmatic diploid examples, one from a diploid Prostate Adenocarcinoma (PRAD), and one from an ovarian (OV) cancer with a highly re-arranged genome. Both samples could be consistent with a potential hopeful monster karyotype. For the former, `TickTack` infers that these copy gains have been acquired in three distinct instants, corresponding to early, mid and late pseudotimes (*τ*_1_ ≈ 0.18, *τ*_2_ ≈ 0.57, *τ*_3_ ≈ 0.94) with 33, 31 and 42 segments in each cluster. In the OV sample, the level of aneuploidy was higher and we identified two clusters of gains, with 94 co-occurring events (92% of total, in 16 chromosomes) at *τ*_1_ ≈ 0.06. In the second cluster, instead, we timed a late trisomy gain of chromosome 7 (*τ*_2_ ≈ 0.84). We show other cases with complex genomes, including WGD ones, in Supplementary Fig.S4-S5.

**Fig. 3.**
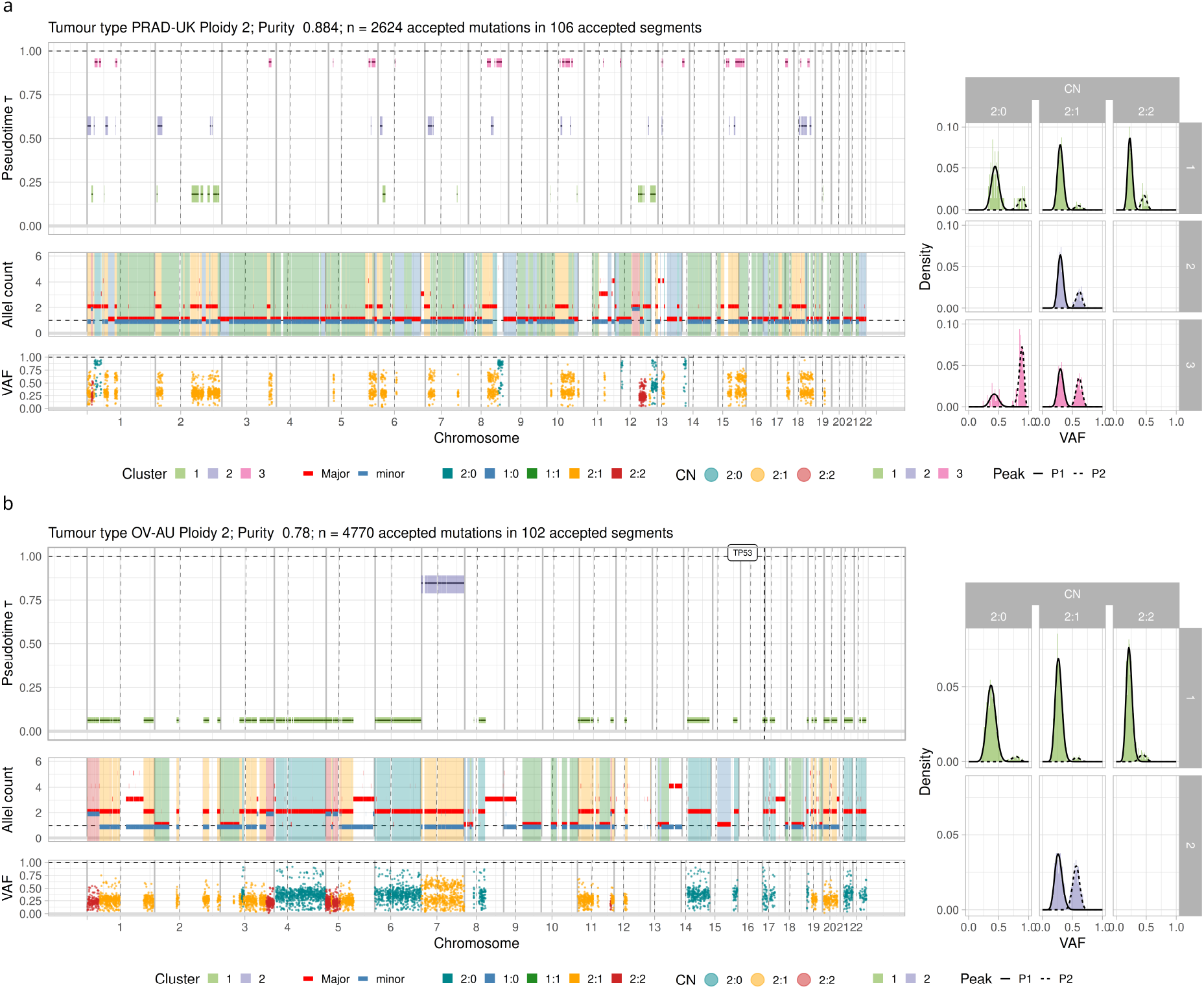
PCAWG inferences potentially compatible with hopeful monsters. **(a)** Prostate adenocarcinoma (PRAD) with diploid genome (ploidy ~ 2). identified three clusters of copy gains (*τ*_1_ ~ 0.18, ~ 0.57, *τ*_3_ ~ 0.94), suggesting three major evolutionary leaps. Most alterations are trisomies, as shown in the allele-specific copy number profile and genome-wide VAFs **(b)** Ovarian (OV) cancer sample with diploid genome and two main clusters: early multiple copy gains at *τ*_1_ ~ 0.06, and a late trisomy of chromosome 7 at *τ*_2_ ~ 0.84.

We attempted to identify hopeful monsters (HM) through clustering co-occurring copy gains. We defined HMs from non-WGD samples with clusters of gains mapped to more than 50% of the chromosomes (Fig.4a and Supplementary Fig.S6). With this definition (Fig.4b), the proportion of HMs across various tumour types was variable, with the highest prevalence observed in ESAD (27.5%), OV (22.8%) and BOCA (20.3%) (Supplementary Fig.S7). Saltational evolutionary patterns involving the simultaneous acquisition of multiple CNAs can be attributed to several biological mechanisms, including chromothripsis, chromoplexy, break-induced replication, breakage-fusion-bridge cycles, and homologous recombination repair defects [37, 38]. Identifying HMs with our model could help map these mechanisms to specific cancer types or mutations (Fig.4c). For example, we observed TP53 mutations to be significantly associated with HMs in ESAD (75% vs 42.8% non-HM). In BRCA, where homologous recombination deficiency plays a crucial role [37], BRCA1 mutations were present in 34.3% of HM and 3.61% of non-HM, while BRCA2 mutations were present in 17.2% of HM and 1.2% of non-HM (Supplementary Fig.S8). Notably, we also observed a difference in the distribution of mutations on RAD51, a key gene involved in the pathway of homologous recombination, in OV (50% of HM vs. 12.9% of non-HM) and PBCA (100% of HM vs. 29.5% of non-HM)(Supplementary Fig.S8).

**Fig. 4.**
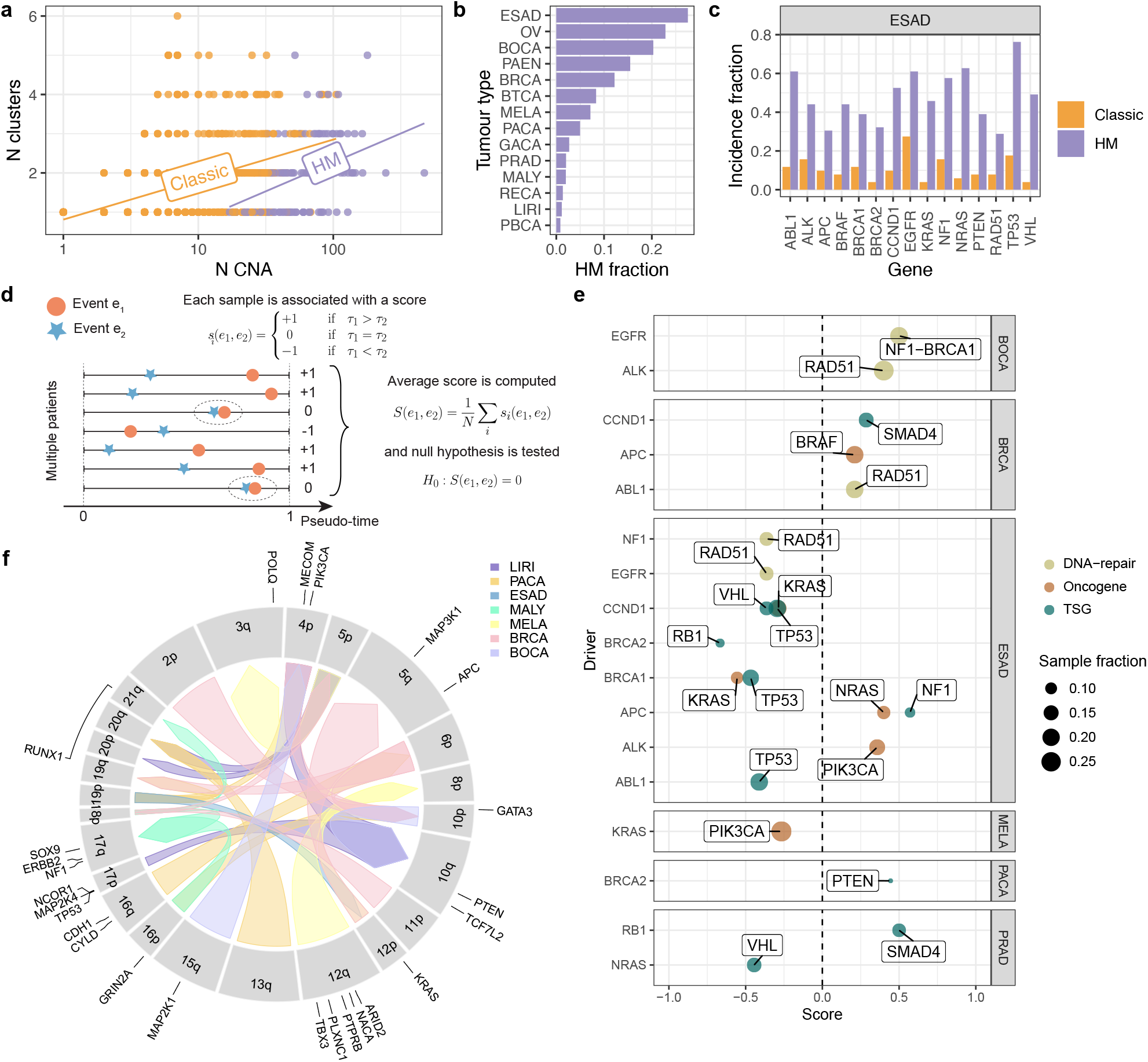
Hopeful monsters and recurrent temporal patterns in cancer evolution. **(a)** Pan-cancer PCAWG linear regression between the number of copy gains and the number of clusters inferred by `TickTack`. We classified samples with potential records of hopeful monsters (HM) searching for clusters with segments mapped to more than 50% of chromosomes (others are Classic). **b)** Distribution of HM samples across cancer types in PCAWG. The highest prevalence is observed in esophageal adenocarcinoma (ESAD, 27.5%), ovarian cancer (OV, 22.8%), and bone cancer (BOCA, 20.3%). **c)** Enrichment of driver gene mutations in HMs samples, determined with p-values of Fisher’s exact test with Benjamini-Hochberg correction. **d)** Schematic of the scoring method used to identify recurrent temporal patterns. For each pair of events (*e*_1_, *e*_2_), a score of +1 or −1 and o is assigned when *e*_1_ occurs after or before *e*_2_; 0 is for co-occurrent gains. The final score is averaged across all samples where both events are observed. **e)** Significant temporal relationships between driver gene alterations, including SMAD4 following RB1 in PRAD and TP53 preceding CCND1, BRCA1, and ABL1 in ESAD (*p* ≤ .05, t-test). **f)** Circos plot showing recurrent temporal patterns of arm-level copy gains across cancer types. Arrows indicate temporal relationships between events, with the arrow direction going from the earlier event to the later event. As an example, it includes early 17p loss (TP53 locus) preceding 10q gain (PTEN locus) in LIRI (*p* ≤ .05, t-test).

Finally, we searched for repeated evolutionary trajectories [39] by examining recurrent sequences of copy gain across various cancer types. To identify these trajectories, we computed a score to determine the temporal consistency of timed gains. For a given pair of events (*e*_1_, *e*_2_) co-occurring in *N* samples with pseudo-times ***τ***_1_, ***τ***_2_, we computed a vector of scores **s**, whose elements are defined as

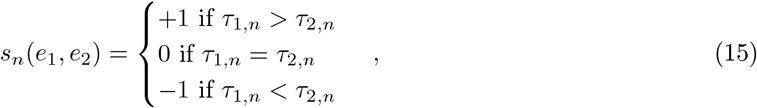

leading to a final score from the average of **s**. This score is negative when the first event is frequently timed before the second and positive otherwise. We then assigned a p-value to each score, testing for significant differences at *α* = 0.05 (t-test against 0; Fig.4d), and classified the events of interest into two categories: focal copy gains affecting known driver genes, versus arm-level copy gains. Our analysis revealed recurrent temporal patterns in driver mutations, suggesting structured progression dynamics in tumour evolution. For instance, in prostate carcinoma (PRAD), SMAD4 alterations appeared to follow RB1 mutations (Fig.4e, Supplementary Fig.S9), a finding supported by previous studies [40, 41]. In esophageal adenocarcinoma (ESAD), TP53 mutations preceded three driver mutations in CCND1, BRCA1, and ABL1. While no definitive evidence confirms that TP53 mutations always occur before these specific drivers, strong evidence suggests that TP53 mutations represent an early event in ESAD, occurring during the transition from normal epithelium to dysplasia [42]. Regarding arm-level copy gains, we observed alterations in loci of well-known cancer genes and patterns in less studied chromosomes. We found that CNLOH of the 17p arm (67% of the cases), where TP53 is located, occurred consistently early in LIRI, where it preceded the amplification of 10q (tetrasomy in 52% of samples), where PTEN is found. The 12p arm, the locus of KRAS, was mostly amplified (single-copy gain, 65% of cases) and constituted an early event in BRCA, where it preceded the copy gain of 5q, but was a late event in ESAD, where it followed the copy gain of 19p (Fig.4f).

## 5 Conclusion

In this work, we have introduced `TickTack`, a hierarchical Bayesian mixture model for reconstructing accurate temporal orders of cancer copy gains while detecting their co-occurrence patterns. Our approach enables understanding complex aneuploidy patterns, including elusive concepts like Goldsmith’s hopeful monsters [11].

Using WGS data from PCAWG, we identified distinct evolutionary patterns across tumour types. By pinpointing clusters of co-occurring events, we attempted to assess the role of saltational evolution in cancer development. While our analysis is still preliminary and depends on several factors (e.g., cutoffs for classification, input genome fragmentation etc.), it is the first automatic attempt at classifying complex evolutionary events recorded in WGS data. Following earlier works [39], our analysis also highlighted recurrent focal driver amplifications and arm-level gains as surrogate patterns of repeated evolution. The ability to distinguish hopeful monsters and uncover these evolutionary patterns holds significant clinical potential, as it unravels targetable complex chromosomal rearrangement mechanisms and enables advanced evolution-based patient stratification techniques. Either way, these efforts could contribute to facilitating personalized treatment strategies by offering new insights into the relationship between cancer drivers and chromosomal instability.

While `TickTack` represents a significant advancement in copy number timing, future work needs to improve the type of supported copy number events, relax simplifying model assumptions (e.g., constant mutation rates) and eventually map genomic events on the actual clinical timeline of a patient. Nonetheless, our findings contribute to the growing body of research leveraging computational and statistical methods to advance cancer biology, offering a flexible framework for studying tumorigenesis, aneuploidy and the evolutionary trajectories of cancer genomes.

## Acknowledgments

The research leading to these results has received funding from AIRC under MFAG 2020 - ID. 24913 project – P.I. Caravagna Giulio. We acknowledge financial support under the National Recovery and Resilience Plan (NRRP), Mission 4, Component 2, Investment 1.1, Call for tender No. 1409 published on 14.9.2022 by the Italian Ministry of University and Research (MUR), funded by the European Union – NextGenerationEU– CUP J53D23015060001.

